# Abnormal autonomic regulation of heart rate underlies age-related and sex dependent cardiac dysfunction in the small primate Microcebus murinus

**DOI:** 10.64898/2025.12.01.691505

**Authors:** Manon Marrot, Pierre Sicard, Yasmine Colombani, Mélanie Faure, Joel Cuoq, Faustine Hugon, Valentin Roger, Antoine Goumard, Nadine Mestre-Frances, Stéphanie Barrère-Lemaire, Matteo E. Mangoni, Angelo G. Torrente

## Abstract

Ageing is a major risk factor for cardiovascular disease associated with the decline of maximal heart rate (HR) and HR variability (HRV). Such a decline could reduce cardiac adaptability to physiological needs and increase susceptibility to arrhythmias. The small primate Grey Mouse Lemur (*Microcebus murinus*), is a valuable model of ageing because it spontaneously develops neurodegenerative diseases. Moreover, in a lifetime it generates about three billion heartbeats as humans. Thus, it could also represent a new effective model to study cardiac ageing.

We investigated cardiac responses to handling stress in young (1-5 years) and aged (6-12 years) lemurs of both sexes by using a mini-Holter for electrocardiogram (ECG) recordings. Moreover, we measured urinary catecholamine, the effect of pharmacological blockade of the autonomic modulation and cardiac contractility.

Aged lemurs, particularly aged males, displayed slower HR and abnormal HRV under stress, delayed recovery of cardiac rhythm following stress and a dissociation between HR and HRV metrics. They also exhibited reduced adrenaline levels, a blunted response to pharmacological vagal blockade and diastolic dysfunction with preserved ejection fraction.

These findings indicate that ageing, especially in male lemurs, impairs cardiac adaptation and recovery from stress through disrupted autonomic balance and sympathetic incompetence.

## 1 Introduction

According to the *World Health Organization* (WHO), the progressive increase in the number of people aged over 60 will constitute a major societal problem in future decades. This demographic shift in our societies will substantially increase the incidence of cardiovascular diseases, which are already the leading cause of death worldwide^1^. Understanding the physiological mechanisms behind the progressive impairment of cardiac function with ageing is therefore essential to develop preventive strategies against the onset of cardiovascular disease in the elderly populations. In addition, sex-related differences influence cardiac physiology and could take part in the ageing process. Indeed, women generally exhibit higher resting HR and parasympathetic modulation^2–4^, while men are characterized by lower responsiveness to adrenergic stimulation^3^. Investigating how these differences contribute to ageing is thus important to prevent cardiac degeneration as reliably and as early in both sexes.

The intrinsic alteration of heart activity with ageing, due to fibrosis and pacemaker dysfunction, has been largely studied in recent years^5^. However, impaired autonomic modulation during ageing may also play a central role in reducing the heart’s responsiveness to body needs and physiological stressors. Indeed, cardiac and brain functions are closely interlinked and age-related neurodegeneration may contribute to alter the autonomic modulation of the heart in aged people^6^. In addition, the decrease of intrinsic resilience with age could increase frailty and decrease the ability to recover from a deviation of normal condition caused by various stress factors.

To explore the mechanism underlying cardiac decline and stress responsiveness with aging, we examined both the intrinsic generation and autonomic modulation of HR in male and female of Grey Mouse Lemurs (GMLs, *Microcebus murinus*). This small primate shares genetic and physiological similarity with humans and has been already used as a model to study age-related decline of physical^7^ and neurological^8^ functions. Indeed, unlike rodents and similar to humans, GML spontaneously develops neurodegenerative disorders similar to Alzheimer’s disease^9^, which could influence the autonomic regulation of cardiac functions.

In addition, we previously demonstrated that the GML heart generates approximately three billion beats in a lifetime, a number of heartbeats comparable to humans and roughly three times greater than mice and other mammals used in biomedical research^10^. Given this similarity and the previous characterization of neurological ageing process, the GML represents a promising model for comparative studies of cardiac ageing. GMLs also offer practical advantages, including a short reproductive cycle (2 months gestation, 40 days parental care, adulthood at 1 to 2 years)^11^ and an easy adaptation to the condition of animal husbandry^12^. Moreover, the GML lifespan is about 5 years in the wild and up to 12 years in captivity^9,10^ and based on different ageing markers, they are considered adult from 1 to 5 years and aged from 6 to 12 years^9,13^. Moreover, we estimated that 12 years of age in GMLs correspond approximately to 85 years in humans^10^. Unlike the preclinical mouse model, whose genetic parameters are controlled, our semi-wild GML colony exhibits also significant heterogeneity, making it more representative of the human physiological diversity^14^.

Taking advantage of this new animal model for cardiac ageing, we investigated the effects of ageing and sex on heart rate (HR) and HR variability (HRV), by using a custom vest holding a mini-Holter to record electrocardiograms (ECGs). In addition, we assessed the autonomic regulation of GML cardiac activity by quantifying urinary catecholamines and their metabolites and using pharmacological blockade of sympathetic and parasympathetic inputs. Further analysis performed by echocardiography allowed us also to investigate age- and sex-related alterations in cardiac contractility.

## 2 Results

### Aged GMLs show an impaired recover of cardiac activity after handling stress

To assess the impact of ageing on GML cardiac activity, we fitted an ECG vest in animals of different age and both sexes, to analyze HR and HRV (**Fig. 1**). In the following text, we indicated adult animals from 1 to 5 years old as young and animals from 6 to 12 years old as aged. The physical restraint required to apply the ECG vest, although non-invasive, imposed a mild stress condition^15^. We therefore considered ECG data recorded just after restraint as the “stress” condition for HR and HRV measurements and data obtained 3h after release as the basal physiological state “post-stress” for these animals.

**Figure 1.**
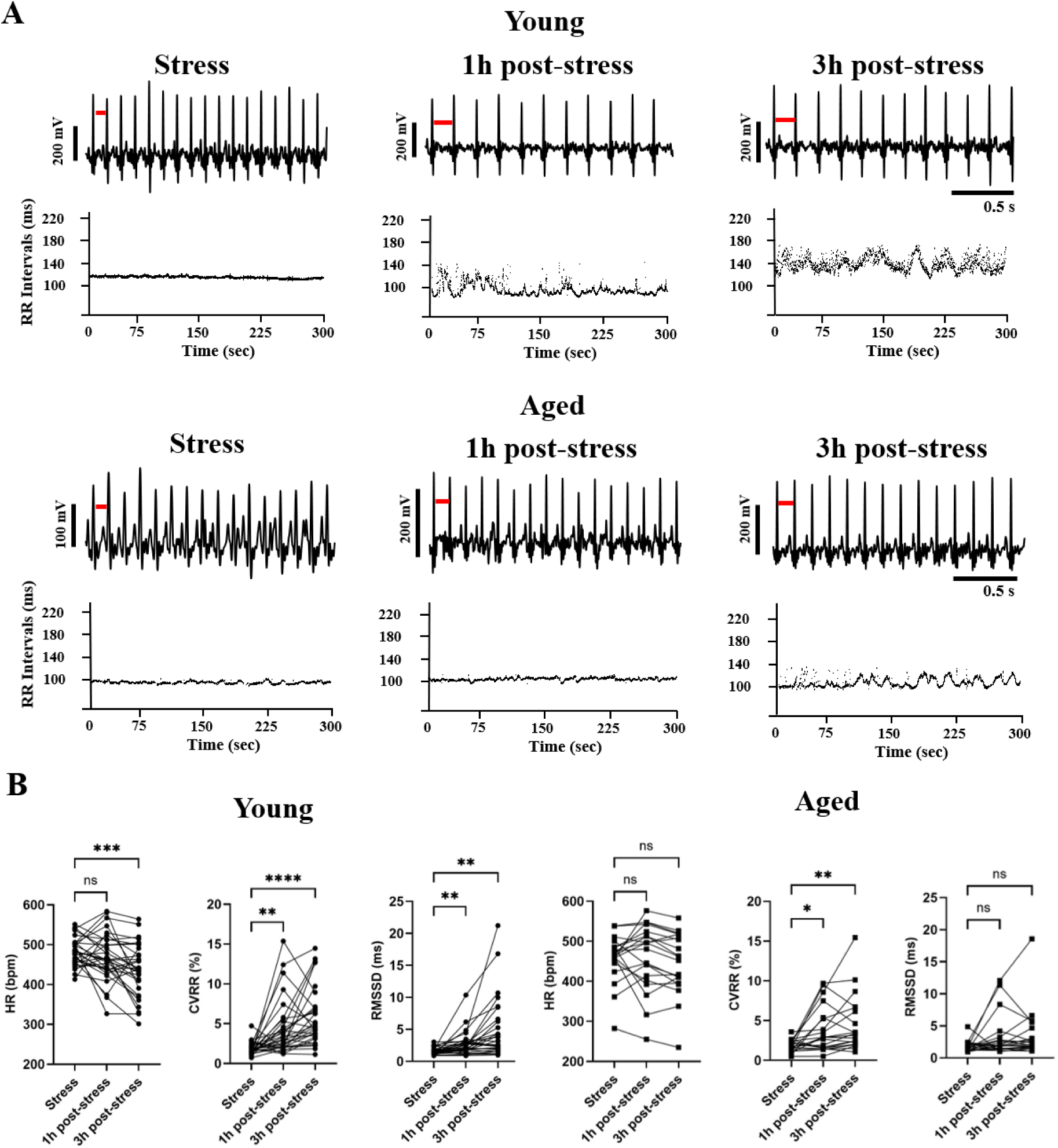
Impaired recover from stress in aged animals. **A.** ECG traces and dot plots of RR intervals in young and aged GMLs recorded under stress, 1h and 3h post-stress. **B.** HR, CVRR and RMSSD in young (n=33) and aged (n=20) GMLs recorded under the condition of **A**. Red dashes highlight RR differences between different stress conditions. *P<0.05, **P<0.01, ***P<0.001, ****P<0.0001 by one-way ANOVA or Friedman test.

Following restraint, in young animals we observed a gradual and significant decrease in HR and increase in two HRV indices, the Coefficient of RR Variability (CVRR), which reflects global HRV and the Root Mean Square of Successive Differences of RR intervals (RMSSD), which measure the beat-to-beat variability (**Fig. 1B**). These results were consistent with the expected normalization of HR and HRV when animals recover from stress.

In contrast, aged GMLs showed no significant differences in HR and RMSSD between stress and post-stress conditions (1h and 3h after restraint; **Fig. 1**). CVRR increased significantly at both 1h and 3h after restraint although to a smaller extent than in young animals (**Fig. 1B**, Suppl. Table 1), suggesting only partial recovery from stress.

Together, these findings indicated that young GMLs normalized their HR and HRV 3h after release, whereas aged GMLs exhibited a prolonged stress effect on cardiac activity that remains detectable 3 h after restraint.

### The HR of aged GML females is less affected by ageing than males

When comparing data from young and aged GMLs, we also observed that aged animals exhibited a lower HR during stress condition, an effect that gradually disappeared 1 and 3 h after release (**Fig. 2A, 2D**).

**Figure 2.**
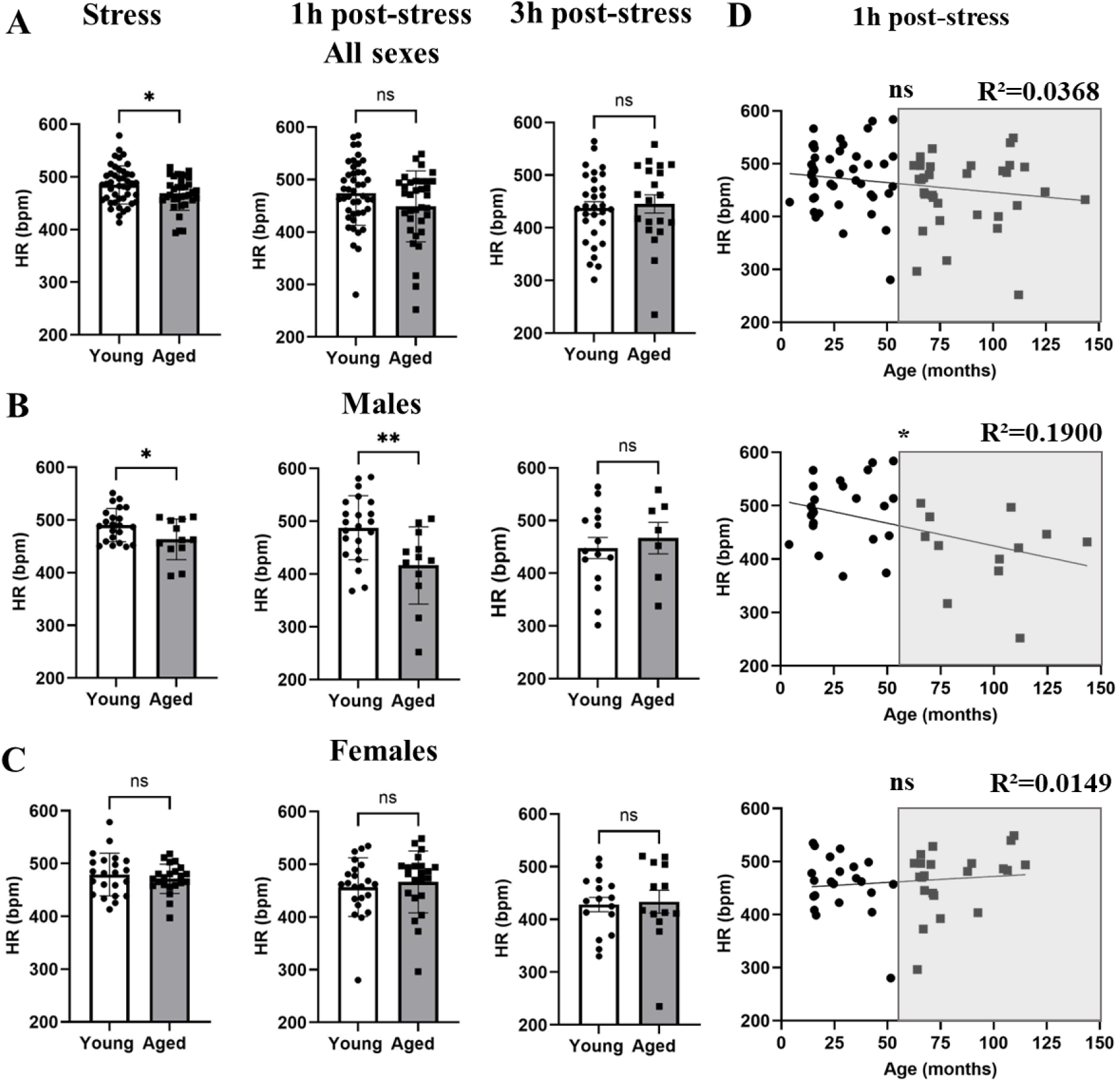
Only the HR of aged GMLs male is affected by ageing. (**A**) HR in young and aged GMLs under stress (young, n=44 and aged, n=34), 1h post-stress (young, n=44 and aged, n=34) and 3h post-stress (young, n=31 and aged, n=20). (**B**) HR in young and aged males of GML under stress (young, n=22 and aged, n=12) 1h post-stress (young, n=22 and aged, n=12) and 3h post-stress (young, n=15 and aged, n=7). (**C**) HR in young and aged females of GML under stress (young, n=22 and aged, n=22), 1h post-stress (young, n=22 and aged, n=22) and 3h post-stress conditions (young, n=16 and aged, n=13). (**D**) Correlation between age and HR 1h post-stress for the same animals reported in (**A**), (**B**) and (**C**) (n=79 for both sexes combined, n=35 in males and n=44 in females). *P<0.05, **P<0.01, by unpaired t-test or Mann-Whitney test and Pearson’s correlation test.

To investigate the influence of sex on cardiac activity across age groups, we conducted a subgroup analysis, separating males and females. In this comparison, we observed a significantly lower HR only in aged males relative to young ones, under stress and 1h after release (**Fig. 2B, 2D**), consistent with our previous findings^10^. Conversely, we did not observe any HR differences between young and aged females regardless of the stress or post stress condition analyzed (**Fig. 2C, 2D**).

### Effect of ageing on HRV in GML

To investigate how ageing influence the autonomic modulation of heart activity, we examined CVRR and RMSSD. These HRV indices have been widely used in small mammals to assess autonomic regulation of the heart during ageing^16–18^. Under stress and 1h and 3h post-stress, we did not find any significant difference of HRV between young and aged GMLs of either sex (**Fig. 3**), excepted for a significantly higher CVRR in aged males under stress (**Fig. 3B**). Because HRV indices generally follow a tendency inversed to HR^19^, this result confirmed the HR decrease that we found in aged GMLs under stress and suggested an impaired cardiac adaptation to stress. In contrast, young and aged females did not differ in any condition or for either HRV indexes (CVRR and RMSSD; **Fig. 3C**), confirming our HR data.

**Figure 3.**
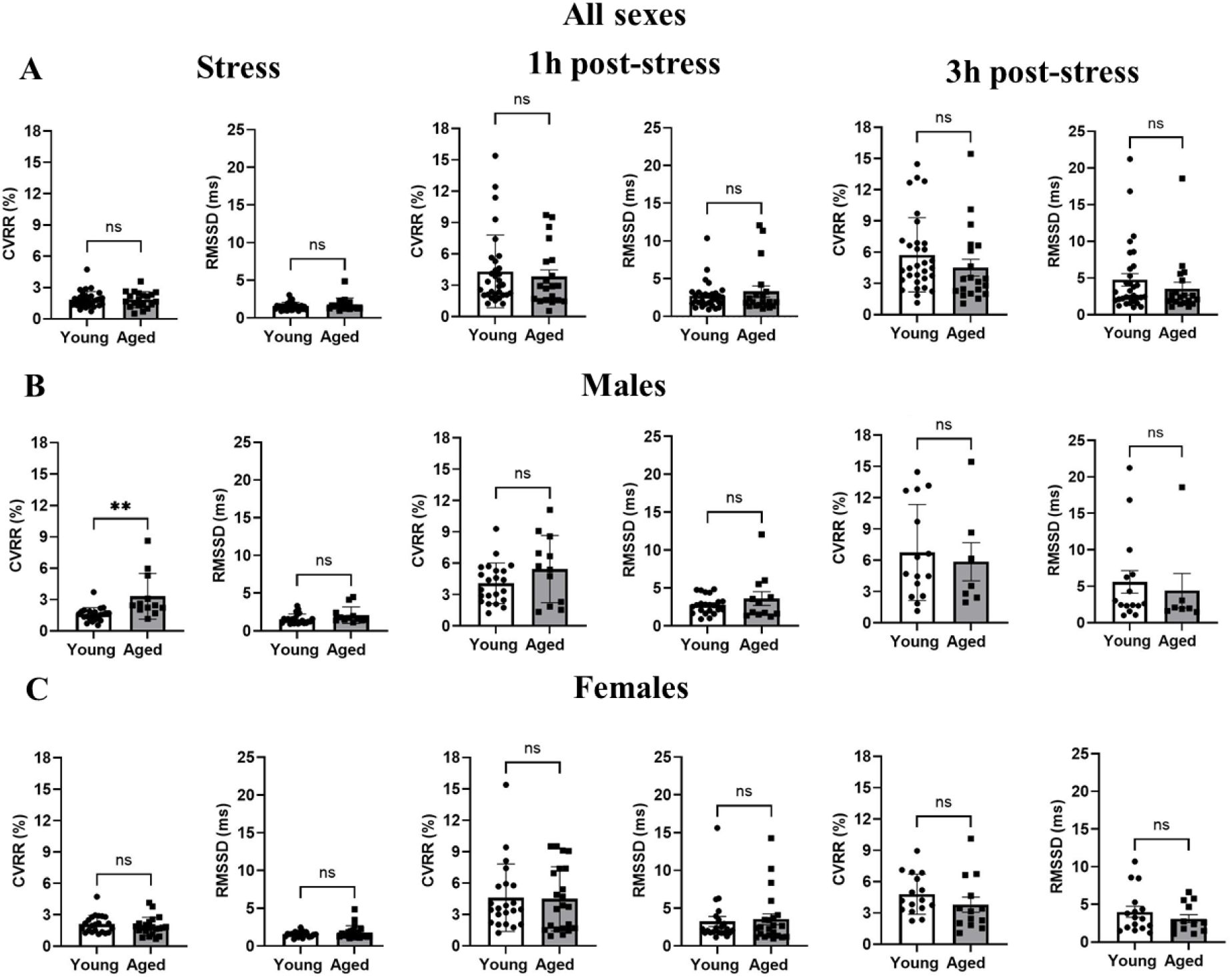
HRV in aged GMLs. (**A**) CVRR and RMSSD recorded for young and aged GMLs under stress (young, n=44 and aged, n=34), 1h post-stress (young, n=44 and aged, n=34) and 3h post-stress (young, n=31 and aged, n=20). (**B**) CVRR and RMSSD recorded only in young and aged males under stress (young, n=22 and aged, n=12), 1h post-stress (young, n=22 and aged, n=12) and 3h post-stress (young, n=15 and aged, n=7). (**C**) CVRR and RMSSD recorded only in young and aged females under stress (young, n=22 and aged, n=22, left panel), 1h post-stress (young, n=22 and aged, n=22, left panel) and 3h post-stress (young, n=16 and aged, n=13, left panel). **P<0.01, by unpaired t-test or Mann-Whitney test.

### Aged GMLs males showed impaired autonomic modulation of cardiac rhythm

Although we did not detect major HRV difference between young and aged animals, the aged group displayed substantial inter-individual variability particularly evident at 1h post-stress, consistent with the heterogeneity of our semi-wild GML colony. To explore this variability, we analyzed individual HRV profiles using CVRR to separate different subgroups. This analysis revealed three distinct HRV subgroups within the aged population, evident also when we separated results obtained from males and females. One subgroup resembled the HRV profile of young animals, whereas the others two showed either low or high HRV compared to young ones (**Fig. 4A**). Therefore, we designed those subgroups as “young-like HRV”, “low HRV” and “high HRV”.

**Figure 4.**
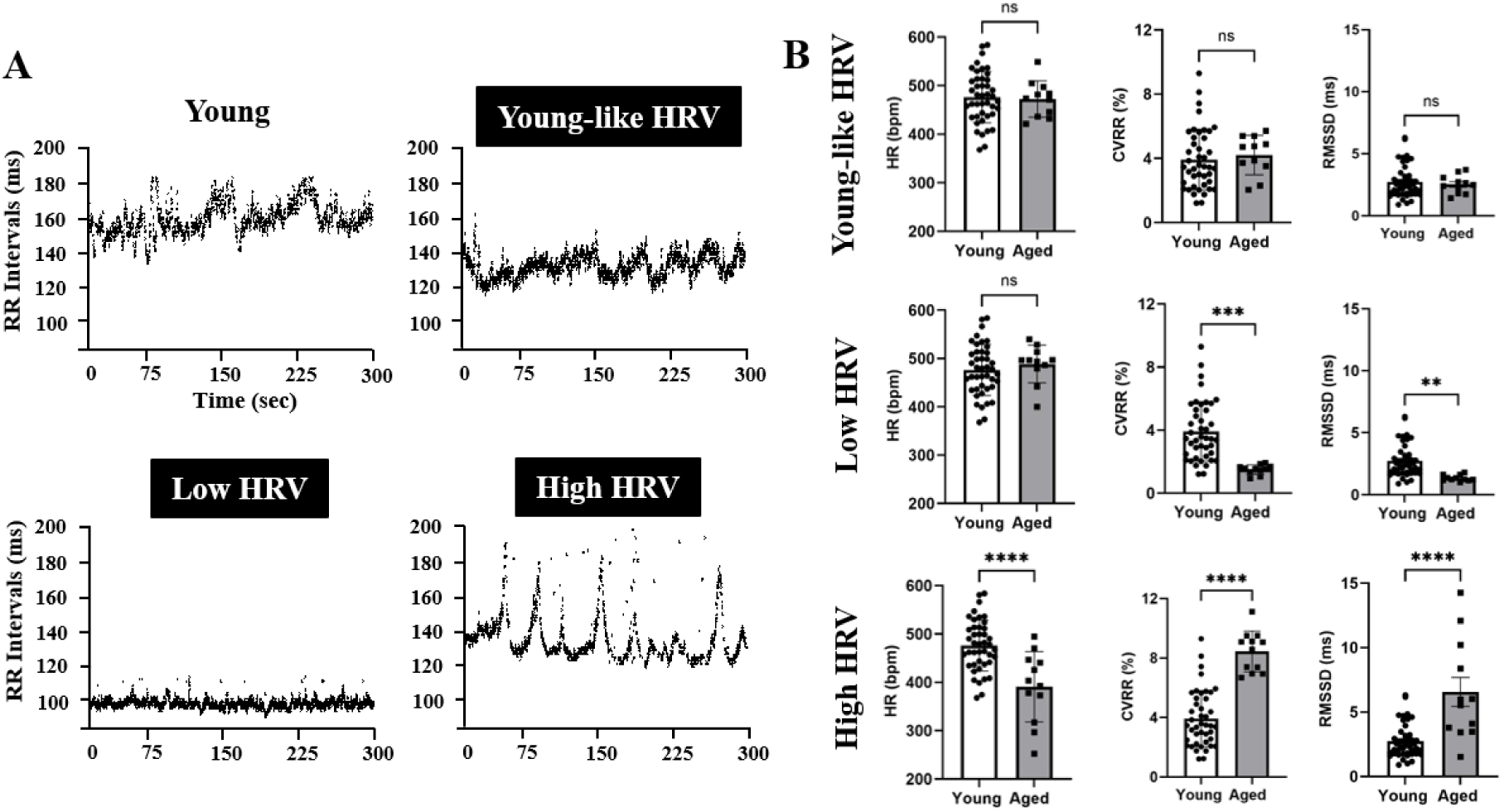
HRV phenotypes discerned in aged GMLs. (**A**) Dot plots of R-R intervals in young and aged animals corresponding to “young-like HRV”, “low HRV” and “high HRV” phenotypes. (**B**) HR, CVRR and RMSSD for the “young-like HRV”, “low HRV” and “high HRV” phenotypes (young, n=43 and aged, n =11). **P<0.01, ***P<0.001, ****P<0.0001, by unpaired t-test or Mann-Whitney test.

We next compared HR and HRV of each of these three subgroups of aged animals with young ones. As expected, “young-like HRV” GMLs did not differed from young ones in HR, CVRR and RMSSD (**Fig. 4B**). In contrast, “low HRV” animals showed significant decrease in CVRR and RMSSD compared to young GMLs, while HR remained unchanged, revealing a dissociation between HR and HRV metrics in this subgroup (**Fig. 4B**). Finally, we noticed a significant decrease in HR and a significant increase in CVRR and RMSSD in “high HRV” aged GMLs in comparison to young animals, consistent with the inverse relationship between HR and HRV^19^ (**Fig. 4B**).

### Effects of pharmacological inhibition of autonomic nervous system input in GMLs

To investigate whether the abnormal HRV profile of aged males might reflect impaired autonomic modulation of cardiac activity, we pharmacologically blocked parasympathetic and sympathetic inputs using atropine and propranolol, respectively. HR recordings following injections of saline solution (NaCl), atropine, propranolol, and atropine + propranolol (A+P), revealed similar responses in young and aged GMLs males (**Fig. 5A to 5D**,) consistent with the predominance of sympathetic tone in small mammals^18^. However, aged animals injected with atropine failed to reach the same HR of young ones (**Fig. 5B**), suggesting chronotropic incompetence.

**Figure 5.**
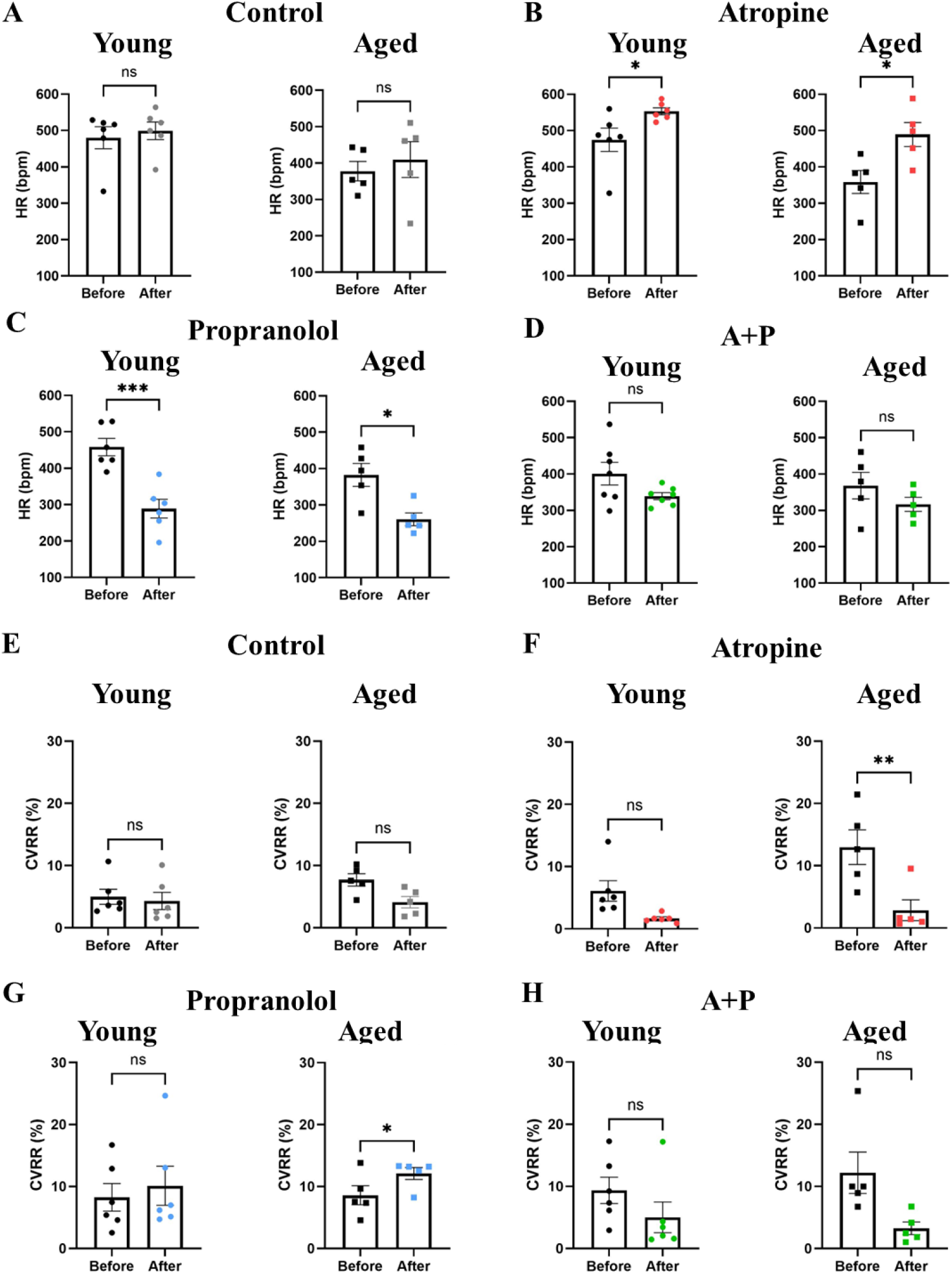
Effects of pharmacological autonomic blockade in young and aged males GMLs. (**A to H**) HR and CVRR in young (n=6) and aged (n=5) before and after the injection with control saline solution (NaCl), atropine (A), propranolol (P) and A+P. *P<0.05, ***P<0.001, by paired t-test.

Moreover, the assessment of global HRV after pharmacological inhibition **(Fig. 5E–H)** further supported this interpretation. Indeed, we noticed that parasympathetic blockade by atropine abolished the increase of CVRR in aged males (**Fig. 5F**). As well, propranolol significantly decreased the HR but increased significantly the CVRR of aged males (**Fig. 5G**), something that we did not observe in young GMLs and could be due to higher vagal tone or intrinsic SAN arrhythmia in aged individuals.

### Aged GMLs show reduced synthesis of catecholamines

To further explore the chronotropic incompetence suggested by the blunted HR increase in aged GMLs during stress or after atropine injection, we evaluated the concentration of catecholamines (CTAs) in urines as a proxy of sympathetic activity. Because restraint during blood collection would immediately increase the concentration of circulant CTAs, we choose to analyze CTAs in urines, which reflects the accumulation of sympathetic activity in the hours before sampling.

According to the reduced HR response to atropine in aged GMLs, we observed a significant decrease of adrenaline (**Fig. 6A**) along with a trend toward decrease in the adrenaline derivate metanephrine (**Fig. 6B**). Conversely, we did not observe difference in noradrenaline (**Fig. 6A**) and its derivate normetanephrine (**Fig. 6B**), as well as in dopamine (**Fig. 6A**) between young and aged GMLs. Similar results were obtained when separating data recorded in males and females (**Fig. 6C, 6D**).

**Figure 6.**
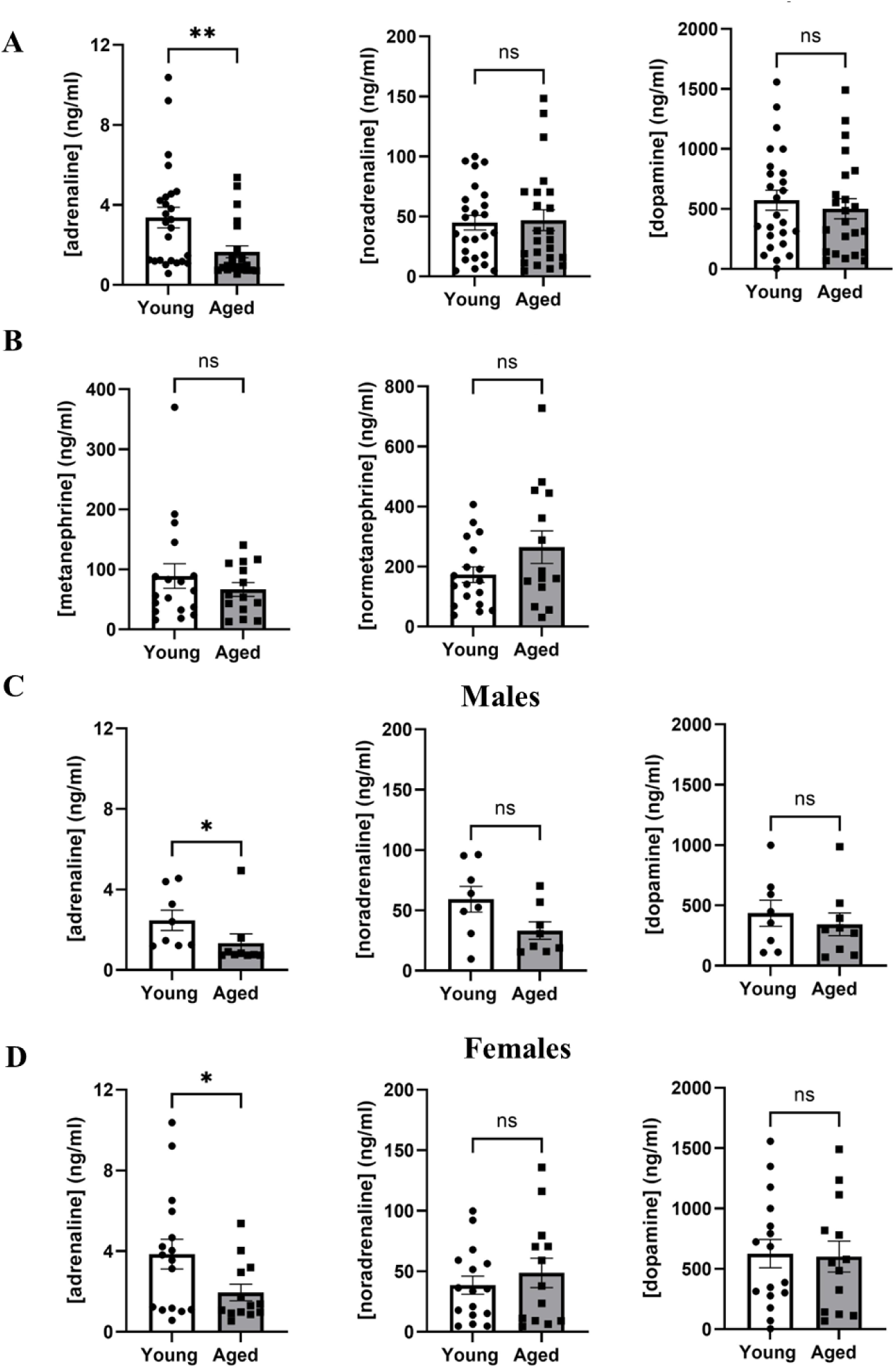
CTA quantification in young and aged GMLs. (**A**) Adrenaline, noradrenaline and dopamine concentrations in the urines of young (n=25) and aged (n=24) GMLs. (**B**) Metanephrine and normetanephrine concentrations in the urine of young (n=18) and aged (n=14) GMLs. (**C**) Adrenaline, noradrenaline and dopamine concentration in the urines of young (n=8) and aged (n=8) GML males. (**D**) Adrenaline, noradrenaline and dopamine concentrations in the urines of young (n=16) and aged (n=13) GML females. *P<0.05, **P<0.01, with an unpaired t-test or Mann-Whitney test.

### Ageing impairs cardiac diastolic function

We then examined the impact of ageing on cardiac function in GML by echocardiography. Conventional systolic parameters showed no difference between young and aged GMLs, as illustrated by similar Left Ventricle Ejection Fraction (LVEF), an index of left ventricular systolic function (**Fig. 7A**). To detect more subtle functional alterations, we applied strain-based echocardiography, which is more sensitive than conventional echocardiography in both clinical practice and preclinical animal models^20,21^. Using this approach, in aged animals we observed a significant reduction in global longitudinal strain (GLS), an early marker of systolic dysfunction (**Fig. 7B**).

**Figure 7.**
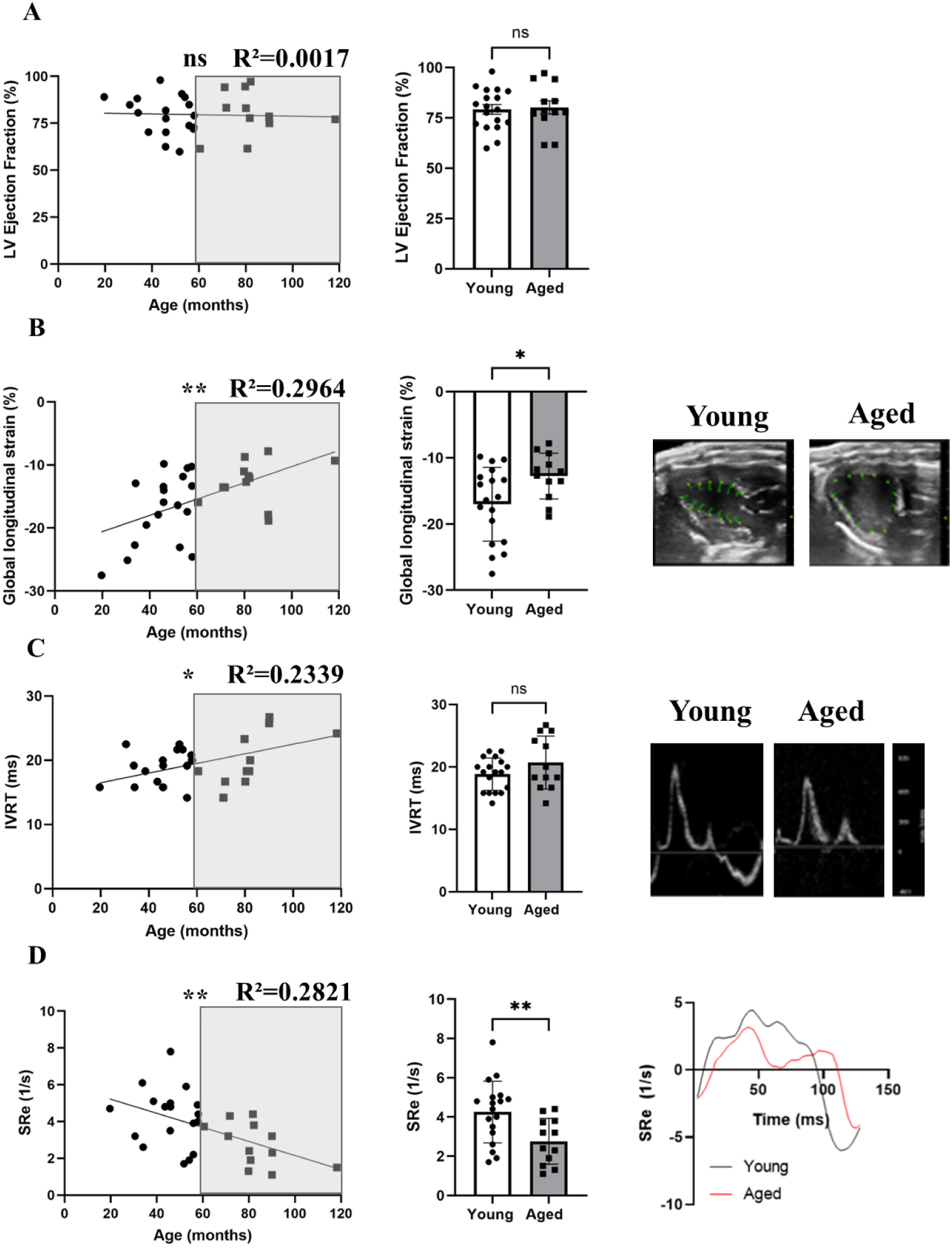
Evaluation of cardiac systolic and diastolic functions in aged animals. (**A**) Correlation between age (months) and LVEF (n=30), in young (n=18) and aged (n=12) GMLs. (**B**) Correlation between age and global longitudinal strain (n=30), in young (n=18) and aged (n=12) GMLs and strain pictures in young and aged GMLs (right panel). (**C**) Correlation between age and IVRT (n=30), in young (n=18) and aged (n=12) GMLs and Doppler pictures in young and aged GMLs (right panels). (**D**) Correlation between age and SRe (n=30), in young (n=18) and aged (n=12) GMLs and traces SRe in young and aged GMLs (right panels). *P<0.05, **P<0.01, with unpaired t-test or Welch’s t test and Pearson’s correlation test.

Moreover, we found that ageing increases the IsoVolumic Relaxation Time (IVRT), a hallmark of diastolic dysfunction (**Fig. 7C**). We also observed a significant decrease in early diastolic Strain Rate (SRe), an index of the strain rate during the early diastole (**Fig. 7D**).

Sex-specific analyses revealed no difference for the LVEF (Suppl, Fig. 1A), but a significant decrease in GLS only in males (Suppl, Fig. 1B). As well we observed an increase in IVRT and a decrease of SRe significant only in females (**Suppl, Fig. 1C and D**). Moreover, both LA area and LV mass significantly increased with ageing (Suppl, Fig. 2A, 2B; Suppl, Fig. 2D, 2E). As expected, in both young and aged GMLs left atrium (LA) area significantly decreases between systole and diastole (Suppl, Fig. 2C).

Altogether, these findings demonstrate an age-related decline in diastolic function in GMLs, similar to humans. As expected, HR in both young and aged animals was lower under anesthesia than in awake animals (Suppl, Table 2), with only a trend toward reduced HR in aged *versus* young GMLs (Suppl, Fig. 3A). We also observed a trend to decrease in body weight with ageing (Suppl, Fig. 3B) and a trend of correlation between HR and body weight (Suppl, Fig. 3C).

## 3 . Discussion

Our study show, for the first time, that ageing modifies the chronotropic and inotropic function of the GMLs’ heart. Indeed, in aged GMLs we observed impaired response and recovery from stress, reduced levels of urinary adrenaline levels, and diastolic dysfunction with preserved ejection fraction. These impairments were more pronounced in GML males, which showed a lower HR than young ones during stress and 1h after stress, suggesting chronotropic incompetence. Aged GML males also exhibited a reduced ability to recover their baseline cardiac activity after stress, suggesting abnormal autonomic modulation^22–24^ and reduced biological resilience, as previously reported in elderly humans^23,24^. In addition, our echocardiography findings indicated both diastolic and early systolic dysfunction with preserved LVEF, similarly to previous observation in humans and mice^19^.

In humans, aged patients are often characterized by the reduction of maximum HR^5^ and decrease of HRV^26^. In aged GMLs, we highlighted that the reduction of HR under stress and 1 h post stress was specific of males and absent in females, confirming previous results showing reduction of sinoatrial node pacemaker activity only in aged men^26,27^.

In aged GMLs males we also observed higher CVRR under stress, a blunted response to vagal blockade with atropine, and reduced levels of adrenaline in the urine, consistent with the significant decrease in epinephrine reported in the urine of age men^28^. Taken together, these results suggest impaired sympathetic response in aged GML males, and a possible reduction in acetylcholine-activated potassium current (I_K,ACh_). In contrast, noradrenaline concentrations did not differ between young and aged GMLs, consistent with studies reporting unchanged urinary noradrenaline concentration in aged men and women^28^, although other authors reported age-related increase in the concentration of noradrenaline in the plasma and urine of patients^29^.

We next investigated the effect of ageing on myocardial function of GMLs. Many studies showed that LV systolic function remains largely preserved with age, while LV diastolic function declines^30^. In our study, we first explored cardiac systolic function, finding no difference in LVEF between young and aged GMLs, confirming data on mice^31^ and humans^32^. On the other hand, although LVEF was not significantly different between young and aged GMLs, GLS was significantly reduced in aged animals, suggesting systolic dysfunction during ageing. Moreover, when separating by sex, aged GMLs males showed impaired systolic function while aged females showed heart failure with preserved ejection fraction (HFpEF), similar to the pattern reported in the “RELAX trial”^33^. These results also align with a previous study done in mice showing higher systolic dysfunction in aged males *versus* female^31^.

Finally, we evaluate diastolic functions, associated with LV remodeling, and in particular with the decline of diastolic function and LA dilatation in aged patients^30,34^. Our results confirmed age-related diastolic dysfunction with an increase in IVRT and a decrease in the rate of early cardiac deformation during diastole. These data are accompanied by an enlargement of left atria with ageing, similar to what has been described for elderly patients^30^ and confirm our previous results on atrial size of aged GMLs^10^. These left atria enlargement in aged patients are associated with several cardiovascular diseases^35^ and could promotes the development of atrial fibrillation^36^.

## 4 . Conclusion

Our results demonstrate that ageing leads to significant alterations in cardiac and autonomic function in GMLs, with marked sex-dependent differences. Aged male showed the most pronounced impairments, including loss of HR adaptation to stress, which may be due to a defective β-adrenergic response to increase HR upon stress, an abnormal vagal tone unable to slowdown HR after stress and a reduction in the SAN response to acetylcholine. Associated with disruption of HR regulation, we also observed cardiac diastolic dysfunction indicative of heart failure and impaired systolic function in males.

Overall, GMLs recapitulate key age-related cardiovascular features observed in humans, supporting their value as a translational model to investigate mechanisms underlying sex-specific cardiac ageing.

## Methods

### Animals

Males and females of GML, were used and subdivided into two age groups: young (1 to 5 years) and aged (6 to 12 years). GMLs were hosted on the colony of the University of Montpellier (license 34-05-026-FS, DDPP, Hérault, France). The experimental protocols involving a non-human primate model have been approved by the French Ministry of Research (APAFiS #43154) and the CEEA-LR Ethics Committee (Committee n°36). GMLs were housed in big cages (1x1x1 m) equipped with enrichment (wooden branches for climbing activities), as well as wooden sleeping boxes imitating the animals’ natural sleeping sites. Room temperature and humidity were maintained at 25 - 27°C and 55 - 65%, respectively. The daily cycles of artificial light and darkness were set to change at 12 pm (light turned off) and were adjusted according to season and to mimic the alternation of day/night, with periods of 6 months of long days similar to summer (14 h of light and 10 h of darkness, noted 14:10) and 6 months of short days similar to winter (10 h of light and 14 h of darkness, noted 10:14). Animals had access to water and food *ad libitum*. They were fed with fresh fruit (apple, orange), applesauce, mealworms and a feed mix (unsweetened condensed milk powder, water, feed supplement, Blédine®, three raw eggs, cottage cheese).

### Electrocardiograms recordings in the conscious animals

ECGs recordings were obtained using a non-invasive external mini-Holter (uECG, Ultimate robotics, Kiev, Ukraine) and electrodes (Asept InMed, Quint-Fonsegrives, France) during the resting period (light time). Animals were shaved on the back with electric shaver, in order to facilitate contact with gel electrodes. To use the ECG device, we designed and homemade vests that held the mini-Holter on the animal’s back. Raw ECG signals collected by the uECG were then analyzed by LabChart pro v8.1.20 (ADInstruments, Dunedin, New Zealand). HR and parameters commonly used to study HRV^37^, such as the Coefficient of Variability of RR intervals (CVRR) and the Root Mean Square of the Successive Differences of RR intervals (RMSSD) were measured to assess cardiac activity and autonomic modulation in GMLs. Because the Standard Deviation of RR intervals (StDRR) is strongly dependent from the mean and the mean RR is lower in aged GMLs compared to young ones, as an HRV index we preferred to use the CVRR, which represent the StDRR normalized to the mean of RR intervals. The animals where then placed in a small transparent cage (37*19*14 cm, L*B*H) for 15 min to adapt to the ECG vest and then were transferred to a normal housing cage for a period of 1 to 3 h, depending on the protocol. HR and HRV analysis were done on 5 min intervals per condition, according to guidelines for HRV analysis in humans^38^.

### Pharmacological injections

Intraperitoneal injections (0.3 mL volume) were performed during the active period (dark time): i) physiological saline (NaCl) solution, ii) 0.5 mg/kg atropine solution (Aguettant, Lyon, France), iii) 5 mg/kg propranolol solution (Sigma-Aldrich, Saint-Louis, Missouri, USA), iv) and 0.5 mg/kg atropine plus 5 mg/kg propranolol solution. These injections were given in winter season, leaving at least 24 h of recovery time between each injection, in order to elimination the molecules from the body of the animal. All molecules were dissolved in physiological saline solution.

### Hormones quantification

We collected urine during the animals’ resting period (light time), to avoid HR changes related to the metabolism of the animals. Almost all GMLs urinate naturally at the time of restraint, as a defense reflex against their predator. These urine collections were systematically carried out before to start the recording of cardiac activity. Urine was stored in a 6 M HCl solution and kept at -20°C protected from light according to the CAT ELISA, MET Urine ELISA protocols (ImmuSmol®, Bordeaux, France).

### Echocardiographic recordings in anesthetized animals and analysis

Echocardiographic recordings were realized in the light period of the day. Animals were measured for body weight before the recording. GMLs were anesthetized with 2-3% isoflurane in an induction chamber and maintained on a controlled heating pad (36-37°C) in dorsal position with 2% isoflurane delivered via 100% O_2_ mask inhalation. Animals were shaved on the chest and abdomen with depilatory cream to facilitate echocardiography imaging. Body temperature and ECG were continuously monitored with rectal probe and electrodes placed on the animals’ limb, respectively. All data were acquired with a high-frequency ultrasound device (Vevo F2, VisualSonics, USA) according to the short- and long-axis view with 2D M-mode imaging and Doppler imaging from an apical 4-chamber view: left ventricle ejection fraction (LVEF), isovolumetric relaxation time (IVRT), early diastolic strain rate (SRE), endocardial global longitudinal strain, left atrium (LA) area were analyzed with Vevo lab software 5.9.0 and Vevostrain 2.0.

### Statistical analysis

Statistical analyses were performed using GraphPad Prism 10.4.2 (GraphPad Software, San Diego, USA). Normality was verified by Shapiro-Wilk test, and on normal data statistical differences were analyzed by t-test or one-way ANOVA. If the normality test was negative, the non-parametric Mann-Whitney test and Friedman test were used. For correlations, Pearson’s test was used. P<0.05 was considered significant. Data were expressed as mean ± standard error of the mean (mean ± SEM). Statistical significance was indicated as * for P<0.05, ** for P<0.01, *** for P<0.001, **** for P<0.0001.

## 5 . Author contributions

MM, PS, YC, MF, VR: acquired data. JC, FH: supervised the animals. MM, YC, VR, AG: analyzed the data. AGT: conceived and designed the research. MM, AGT and SBL: drafted the manuscript. MM, PS and AGT: prepared figures. NMF, SBL, MEM and AGT: made critical revision of the manuscript and handled funding and supervision. All authors contributed to the article and approved the submitted version.

## Acknowledgements

We are thankful to all the members of the RAM-CECEMA animal facility and the IPAM/Biocampus platform for their help and support of our project.

---

**Supplementary Figure 1.**
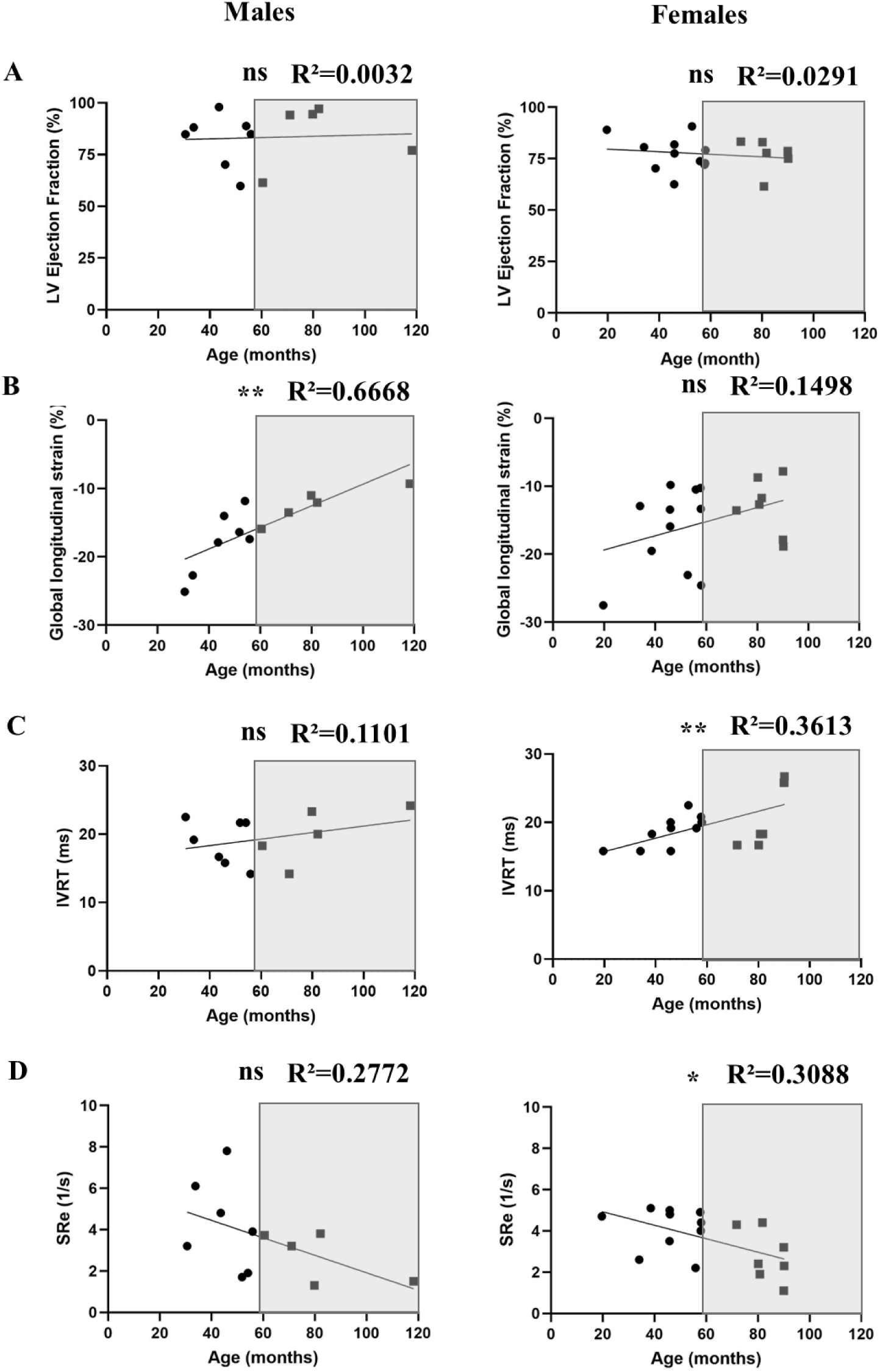
Effect of ageing on cardiac systolic and diastolic functions in male and females of GMLs. (**A, B, C and D**) Correlation between age and LVEF, global longitudinal strain, IVRT and SRe in GMLs males (dot and square indicates young [n=12] and aged animals [n=5], respectively) and females (dot and square indicates young [n=11] and aged animals [n=7], respectively). *P<0.05, **P<0.01, by Pearson’s correlation test.

**Supplementary Figure 2.**
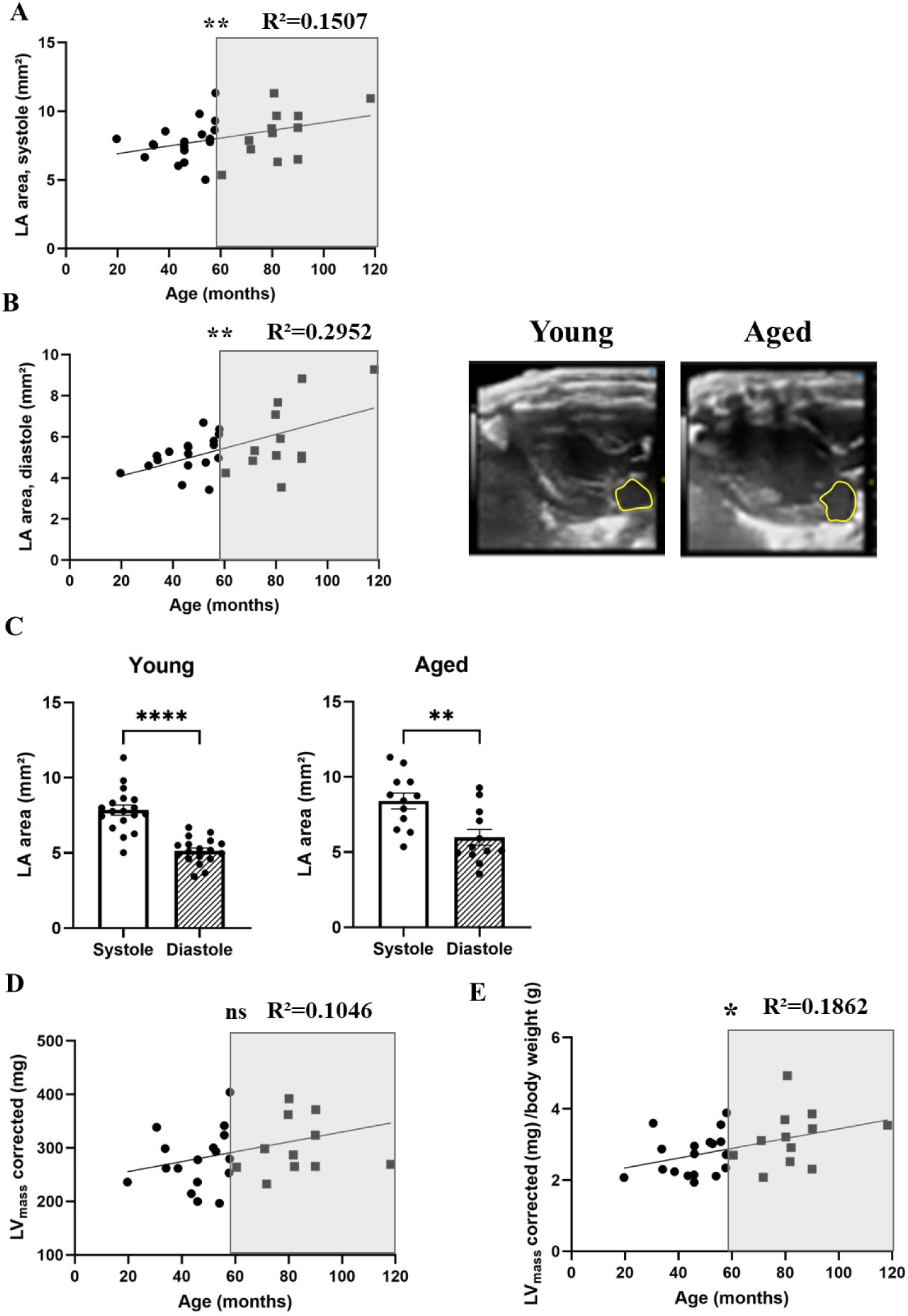
Cardiac systolic and diastolic parameters. (**A and B**) Correlation between age and LA area in systole and diastole (n=30) and LA pictures in young and aged GMLs (right panels). (**C**) Comparison between LA area during systole and diastole in young (n=18) and aged (n=12) GMLs. **(D)** Correlation between age and LV_mass_ (n=30). (**E**) Correlation between age and LV_mass_ corrected for body weight (n=30). *P<0.05, **P<0.01, ****P<0.0001, by unpaired t-test or Welch’s test and Pearson’s correlation test.

**Supplementary Figure 3.**
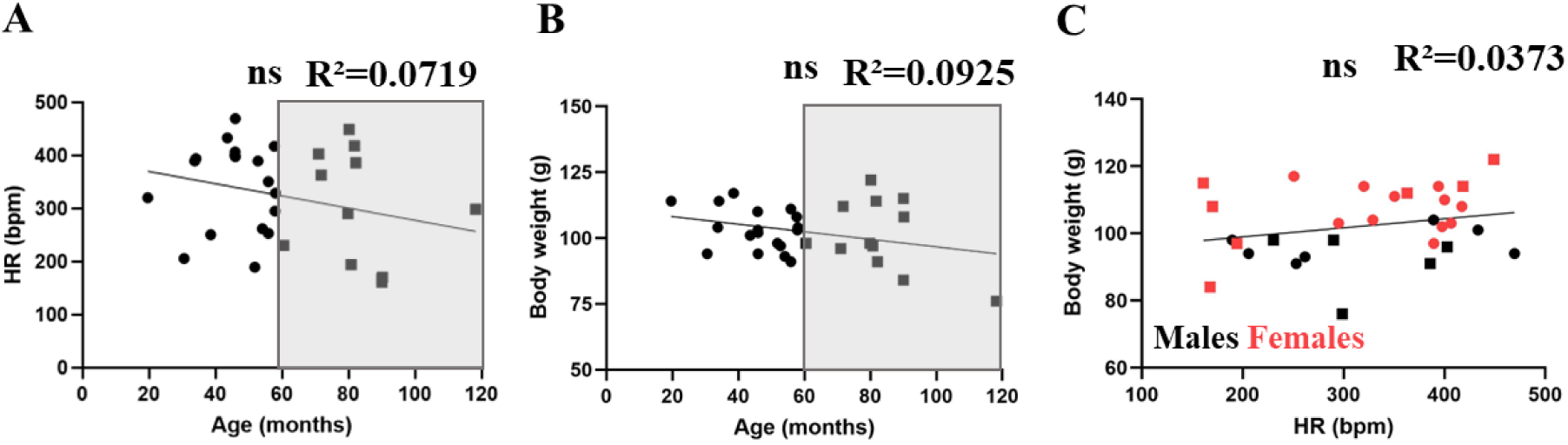
Correlation between HR, body weight and age for animals analyzed by echocardiography. (**A**) Correlation between age and HR recorded under anesthesia in all GMLs (young, n=18 and aged, n=12). (**B**) Correlation between age and body weight (young, n=18 and aged, n=12). (**C**) Correlation between HR and body weight by separating data by sexes (dot and square indicates young [n=18] and aged animals [n=12], respectively). Statistical significance of the correlations was analyzed by Pearson’s correlation test.

**Supplementary Table 1.** Stress effect on HR and HRV. Difference in HR, CVRR and RMSSD expressed as a percentage of change between handling stress and 1h or 3h post-stress. P-values were obtained from the percentage difference between the stress versus post-stress conditions in young and aged GMLs.

|  | HR |  |  | CVRR |  |  | RMSSD |  |  |
| --- | --- | --- | --- | --- | --- | --- | --- | --- | --- |
|  | Young | Aged | p value | Young | Aged | p value | Young | Aged | p value |
| $\Delta$ \|Stress vs. 1h\| | -2.5% | 0.6% | 0.4 | 190.1% | 134.9% | 0.5 | 85.1% | 106.5% | 0.6 |
| $\Delta$ \|Stress vs. 3h\| | -9.1% | 2.9% | 0.1 | 262.6% | 163.1% | 0.1 | 219.7% | 124.3% | 0.2 |

**Supplementary Table 2.**
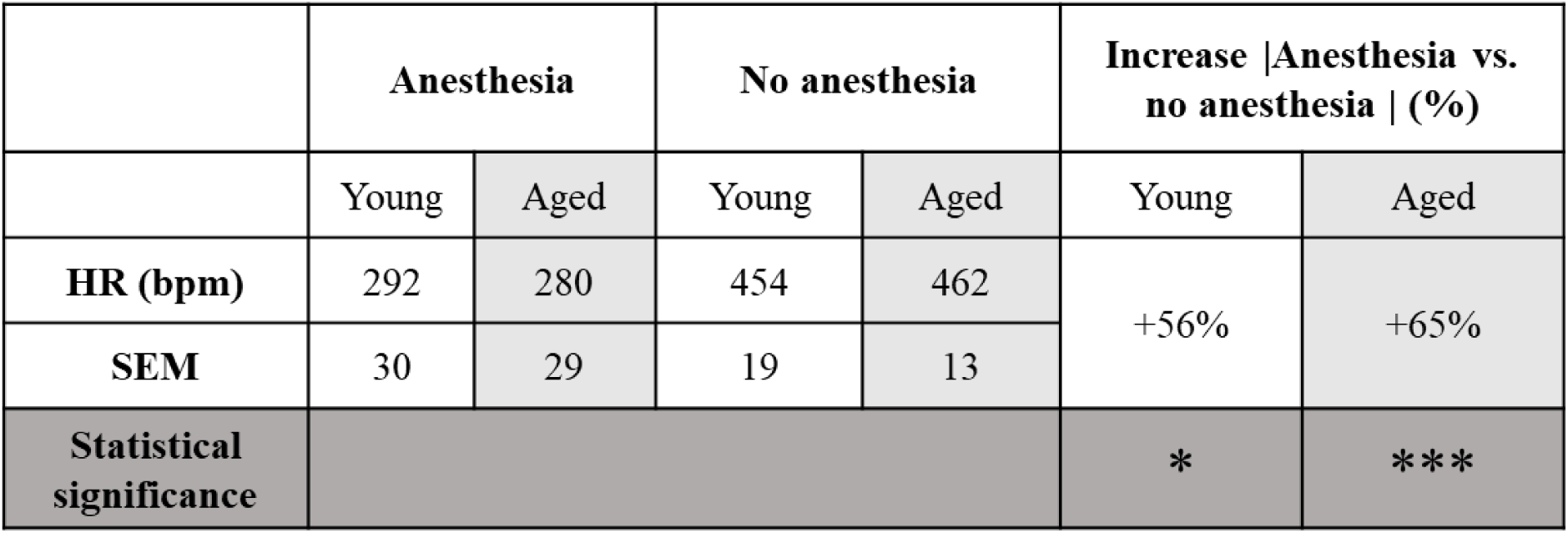
HR recorded with or without anesthesia. HR (bpm) recorded in GMLs under anesthesia or in awake and freely moving animals, in young (n=7) and aged (n=11). Lower line indicates the percentage of increase in HR between anesthetized and not anesthetized young and aged GMLs. *P<0.05, ***P<0.001, with unpaired t-test or non-parametric Welch’s t test.

|  | Anesthesia |  | No anesthesia |  | Increase \|Anesthesia vs. no anesthesia \| (%) |  |
| --- | --- | --- | --- | --- | --- | --- |
|  | Young | Aged | Young | Aged | Young | Aged |
| **HR (bpm)** | 292 | 280 | 454 | 462 | +56% | +65% |
| **SEM** | 30 | 29 | 19 | 13 |  |  |
| **Statistical significance** |  |  |  |  | * | *** |

